# MegaLMM improves genomic predictions in new environments using environmental covariates

**DOI:** 10.1101/2024.03.06.583749

**Authors:** Haixiao Hu, Renaud Rincent, Daniel E. Runcie

**Affiliations:** Department of Plant Sciences, University of California Davis, Davis, CA 95616, USA; Université Paris-Saclay, INRAE, CNRS, AgroParisTech, GQE - Le Moulon, 91190 Gif-sur-Yvette, France

**Keywords:** genotype-by-environment interaction, multivariate linear mixed model, factor analytic model, multi-environment trials, environmental covariates

## Abstract

Multi-environment trials (METs) are crucial for identifying varieties that perform well across a target population of environments (TPE). However, METs are typically too small to sufficiently represent all relevant environment-types, and face challenges from changing environment-types due to climate change. Statistical methods that enable prediction of variety performance for new environments beyond the METs are needed. We recently developed MegaLMM, a statistical model that can leverage hundreds of trials to significantly improve genetic value prediction accuracy within METs. Here, we extend MegaLMM to enable genomic prediction in new environments by learning regressions of latent factor loadings on Environmental Covariates (ECs) across trials. We evaluated the extended MegaLMM using the maize Genome-To-Fields dataset, consisting of 4402 varieties cultivated in 195 trials with 87.1% of phenotypic values missing, and demonstrated its high accuracy in genomic prediction under various breeding scenarios. Furthermore, we showcased MegaLMM’s superiority over univariate GBLUP in predicting trait performance of experimental genotypes in new environments. Finally, we explored the use of higher-dimensional quantitative ECs and discussed when and how detailed environmental data can be leveraged for genomic prediction from METs. We propose that MegaLMM can be applied to plant breeding of diverse crops and different fields of genetics where large-scale linear mixed models are utilized.

## INTRODUCTION

Genotype-by-environment interactions are one of the most difficult challenges faced by plant breeders. Good varieties must maintain performance across a wide range of environments. However, testing every candidate variety in every possible condition within the target population of environments (TPE) is not feasible. Instead, breeders evaluate candidate genotypes in multi-environment trials (METs) covering a moderate number of locations over multiple years. METs can consume a large fraction of a breeding program’s budget. Therefore, making optimal use of data from METs for breeding decisions is critical to the success of plant breeding programs.

Many statistical approaches for modeling data from METs have been developed (as reviewed by Crossa *et al*. (2022)). Historically, most models have taken one of two major approaches: reaction norm models represent the change in the performance across trials as a function of measurable characteristics of those trials, called Environmental Covariates (ECs), while correlated trait models represent the correlation in performances across genotypes between pairs of trials. Examples of reaction norm models include factorial regression (Denis 1988; Piepho *et al*. 1998), the GBLUP-based reaction norm model (Jarquín *et al*. 2014; Ly *et al*. 2018), the Critical Environmental Regressor through Informed Search-Joint Genomic Regression Analysis (CERIS-JGRA) model (Li *et al*. 2021), and models based on Crop Growth Models (*e*.*g*. Technow *et al*. 2015). Examples of correlated trait models include the Additive Main Effect and Multiplicative Interaction (AMMI) approach (Gollob 1968; Zobel *et al*. 1988) and latent factor models (Smith *et al*. 2001; Cullis *et al*. 2014). In their most general forms, reaction norm models and correlated traits models can be mathematically equivalent, and several of these models combine aspects of both approaches. However, each approach has its own computational and statistical advantages. One advantage of the correlated traits approach is that it has the potential to completely characterize the correlation between any pair of trials, while reaction norm models can only learn the components of the correlation that are captured by the ECs utilized to parameterize the reaction norm. Therefore, correlated traits models are expected to be more accurate for the specific trials in the METs. On the other hand, reaction norm models can be used to make predictions in un-measured environments while correlated traits models cannot. Historically, correlated traits models have been less computationally tractable because the number of correlations that must be learned grows quadratically with the number of trials. We recently developed MegaLMM, a computationally and statistically efficient implementation of a multivariate linear mixed model, and demonstrated that it could accurately perform genomic prediction in METs with more than 100 trials, improving predictive ability in nearly every trial relative to univariate approaches (Runcie *et al*. 2021). MegaLMM is a correlated traits model built on a factor analytic (FA) structure, however it lacks of a prediction mechanism to extrapolate genomic values to new environments with unobserved environmental conditions. Extending MegaLMM to make use of ECs would allow it to encompass the benefits of both the correlated traits and reaction norm approaches to modeling data from METs.

The quality of environmental covariates limits the potential of any model to predict genetic values in new environments. Several challenges in developing environmental covariates include: i) there are many environmental variables that impact plant growth and development, including temperature, water availability, soil properties, disease pressure, etc, ii) many of these variables are dynamic, meaning that they change during a growing season on both short and long time scales, and interact with plants differently depending on the growth stages of each plant, iii) environmental factors are collinear and may interact with one another, making statistically identifying causal drivers challenging, iv) some important variables are challenging to measure or are unknown, and v) environmental variables from the growing season are unknown at the time of planting or for future unobserved locations. Because of the high dimensionality of potential ECs, statistical models that use ECs must operate robustly in high-dimensional spaces. There are three common strategies for dealing with high-dimensional ECs: 1) Variable selection. As an example, CERIS-JGRA (Li *et al*. 2021) searches a large set of candidate ECs for the single most useful one and then uses only that one for prediction. 2) Non-linear machine learning such as kernel regression or Deep Learning. The models of Jarquín *et al*. (2014) and Costa-Neto *et al*. (2021) use kernel methods to represent the covariance of environments based on EC distances and performing regressions using these distances. Washburn *et al*. (2021) and Kick *et al*. (2023) utilized deep learning techniques to prioritize ECs with poential agricultural importantce. 3) Crop growth models. Heslot *et al*. (2014) and Technow *et al*. (2015) use biophysical-based models to predict the impact of EC time-series’s across multiple ECs on crop physiology and development. Heslot *et al*. (2014) and Rincent *et al*. (2019) used crop growth models as a form of non-linear dimension reduction to extract a more physiologically relevant set of ECs to use in MET models.

Measuring the success of genotype-environment interaction models from METs is complicated because such models can be used for multiple different tasks in a breeding program. Breeders evaluate samples of genotypes from a reference population of genotypes (RPG) in samples of environments from a TPE (Cooper *et al*. 2021). Genotypes observed in at least one trial of a MET are commonly called “old genotypes” while the remaining genotypes in the RPG are called “new genotypes”. The trials that compose a MET are called “old environments”, while other possible growing environments in the TPE are called “new environments”. Four distinct applications are commonly distinguished: 1) Imputing performances of genotypes in the MET in trials where some genotypes were not grown, for example if a MET is sparse (Burgueño *et al*. 2012); 2) Predicting the relative performances of new genotypes in each of the environmental conditions represented by trials in the MET; 3) Predicting the relative performances of a set of genotypes in new environments, based on their performances in a MET; and 4) Predicting the relative performance of the new genotypes in new environments. The first two applications are statistically easier than the last two, while the fourth is the most difficult because it relies on predicting characteristics of previously unobserved genotypes and environments. MET models should be evaluated in each of these contexts because performance in one context does not guarantee adequate performance in another. The most common computational strategy for evaluating model accuracy is cross validation. Cross validation strategies that simulate each of these applications are termed CV2, CV1, CV3, and CV0, respectively (Burgueño *et al*. 2012; Costa-Neto *et al*. 2021).

Here, we describe an extension to MegaLMM that facilitates the use of ECs to extend genomic predictions to new environments. Our primary objective is to describe the statistical framework of the extended MegaLMM model and evaluate its efficacy in various breeding scenarios. We use a maize hybrid dataset from the Genomes-To-Field (G2F) Initiative (AlKhalifah *et al*. 2018) to demonstrate that MegaLMM can achieve high accuracy in genomic prediction under various breeding conditions. We show that MegaLMM surpassed univariate GBLUP in predicting hybrid performance in new environments partly through its effective use of ECs. Finally, we explore the use of higher-dimensional quantitative ECs and discuss when and how detailed environmental data can be leveraged for genomic prediction from METs.

## RESULTS

### Method Overview

We developed the original MegaLMM model to provide a robust framework for modeling the correlations of genetic values of experimental genotypes across multiple environments. MegaLMM links genetic predictors to phenotypic data using a hierarchical latent factor model that is computationally efficient, yet highly flexible to accommodate different genetic architectures across traits (Runcie *et al*. 2021). MegaLMM decomposes a high-dimensional, but potentially sparsely populated phenotypic matrix (**Y**) into a low-rank factor score matrix (**F**), a low-rank loading matrix (**Λ**), and a residual matrix (**E**) (Figure 1A). Together, the factor matrix and the loading matrix explain the genetic covariation among environments, while the residual matrix accounts for unexplained residual genetic variation, microenvironmental variation, and measurement error unique to each environment. Learning latent factor scores for each individual in the training set allows MegaLMM to predict genetic values of each observed genotype in environments where that genotype was not grown (but other genotypes were, *i*.*e*. the CV2 context) (Figure 1B). Latent regressions of each vector of factors scores (columns of **F**), and each residual vector (columns of **E**) on genetic data from each observed genotype allows MegaLMM to predict genetic values of new genotypes (without any phenotype data in **Y**, *i*.*e*. the CV1 context) in each environment by predicting factor scores **F**_*n*_ and residual values **E**_*n*_ for each new genotype based on inputs of genetic data (Figure 1B). However, the original MegaLMM had no mechanism to link values in **Λ** to external data representing properties of each environment. Therefore, MegaLMM had no mechanism to predict genetic or phenotype values of either observed or unobserved genotypes in new environments.

**Figure 1.**
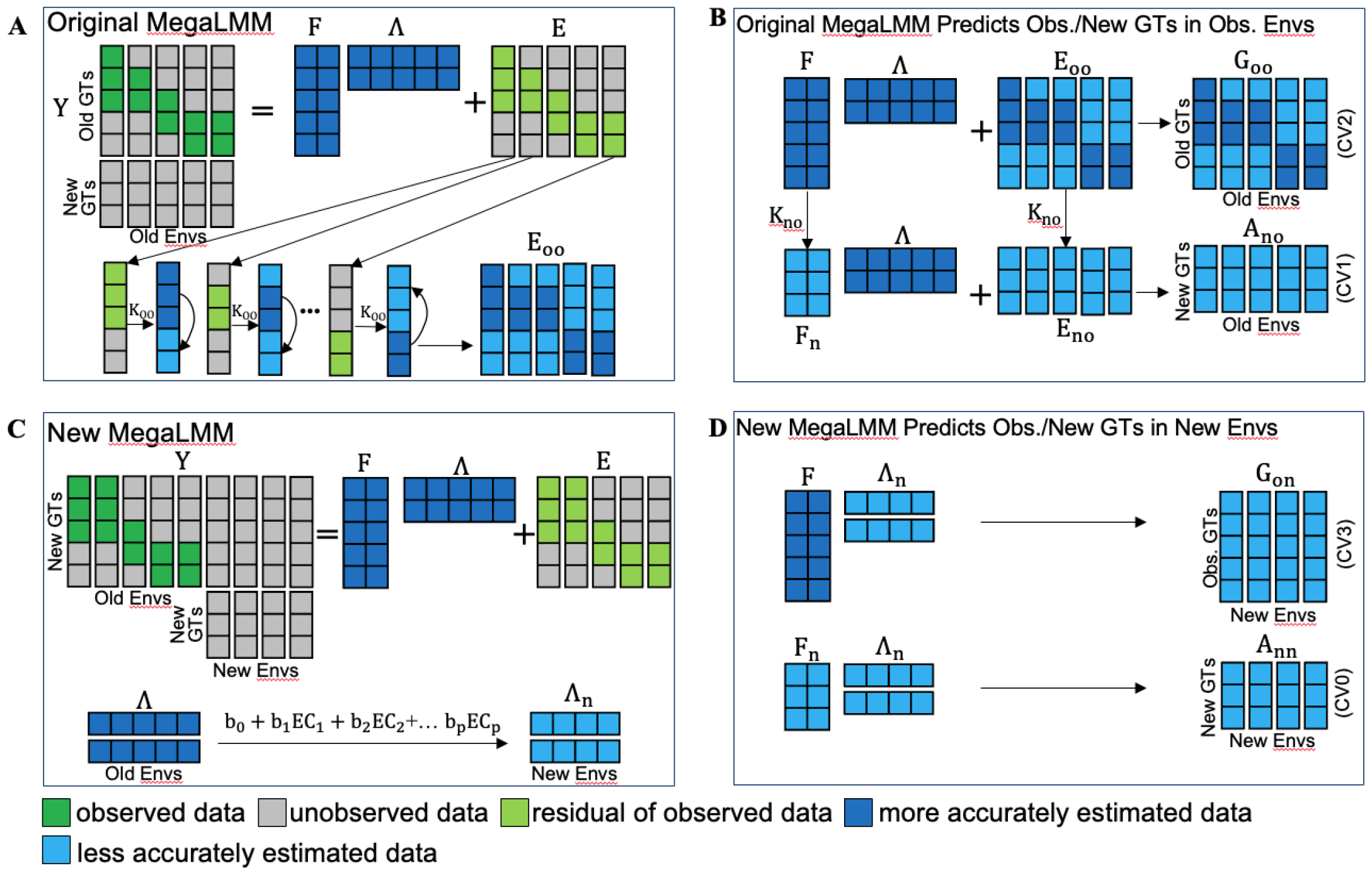
MegaLMM statistical models and their applications for predicting trait performance in experimental genotypes across observed and new environments. (A) Original MegaLMM model architecture. (B) Approach for predicting genetic values of old or new genotypes in old environments using the original MegaLMM. (C) Model architecture of the new MegaLMM model. (D) Approach for predicting genetic values of old or new genotypes in new environments using the new MegaLMM. **Y**: phenotypic matrix consisting of phenotypic values measured on *n* genotypes (rows) in *t* environments (columns). **F**: factor matrix of old genotypes. **F**_*n*_: predicted factor matrix of new genotypes. **Λ**: factor loading matrix for the old environments. **Λ**_*n*_: predicted factor loading matrix for new environments. **K**: additive genomic relationship among old genotypes. **E**: residual trait matrix for observed genotypes in observed environments after accounting for the latent factors. **A**_*on*_: predicted additive genetic values of old genotypes in new environments; **G**_*on*_: predicted total genetic values of old genotypes in new environments. **A**_*no*_: predicted additive genetic values of new genotypes in old environments. **A**_*nn*_: predicted additive genetic values of new genotypes in new environments. **G**_*oo*_: predicted genetic values of old genotypes in old environments. GTs= Genotypes, Envs=Environments, EC=Environment Covariates.

Here, we extend MegaLMM to accept environmental data as predictors of the covariances of genetic values across environments. The extended model keeps all features of the original model, but adds functionality to express the rows of **Λ** as regressions on sets of ECs (Figure 1C). We can then predict genetic values of either old or new genotypes in any new environment that can be characterized by these ECs (Figure 1D). As an intuitive justification for this approach, we consider the variation in trait values in a single environment **y**_*j*_ to be caused by a set of latent characteristics **f**_**1**_ … **f**_**k**_ (such as flowering time, growth rate, and drought tolerance). In a different environment, many of these same latent characteristics will still be important but their relative effects on overall performance may vary. Each row of **Λ** represents the relative importance of a single latent characteristic across environments. For example, if growth rate is similarly important in all environments, values in the corresponding row of **Λ** will be similar. If earlier flowering is beneficial in some environments but detrimental in others, the corresponding row of **Λ** will have some positive and some negative values. Our overall hypothesis is that the variation in these importance weights across environments will be predictable by known characteristics of those environments, including geography, climate, and management. Details of this latent regressions approach are provided in the Methods.

### MegaLMM greatly improves genomic predictions of agronomic traits within experimental trials

We used the G2F maize hybrid dataset (Lima *et al*. 2023), covering the years 2014 to 2021, to evaluate the genomic predictive ability of MegaLMM in its original form and with the enhancements described above. The G2F maize hybrids are crosses between a large set of inbred lines (referred to as P1) and a small set of tester lines (referred to as Tester). We formatted a trait matrix **Y** with P1s as rows and combinations of Tester, location, and year as columns, and filled each value with a least-squares mean estimate of the corresponding hybrid trait values. We subsetted the data to include columns with at least 50 observed trait values, which resulted in data from a total of 12 Testers. The replacement of P1s every two years resulted in a very sparse trait matrix with 87.1% missing values for Grain Yield (Figure 2A). Below, we refer to individual columns of this trait matrix as an “experiment”, signifying trait values from a set of P1s crossed to a single tester and evaluated in a specific location-year combination. In total, our dataset was composed of 1702 P1s, 12 testers, 4402 hybrids, 302 experiments, and 195 trials for Grain Yield (Figure 2A).

**Figure 2.**
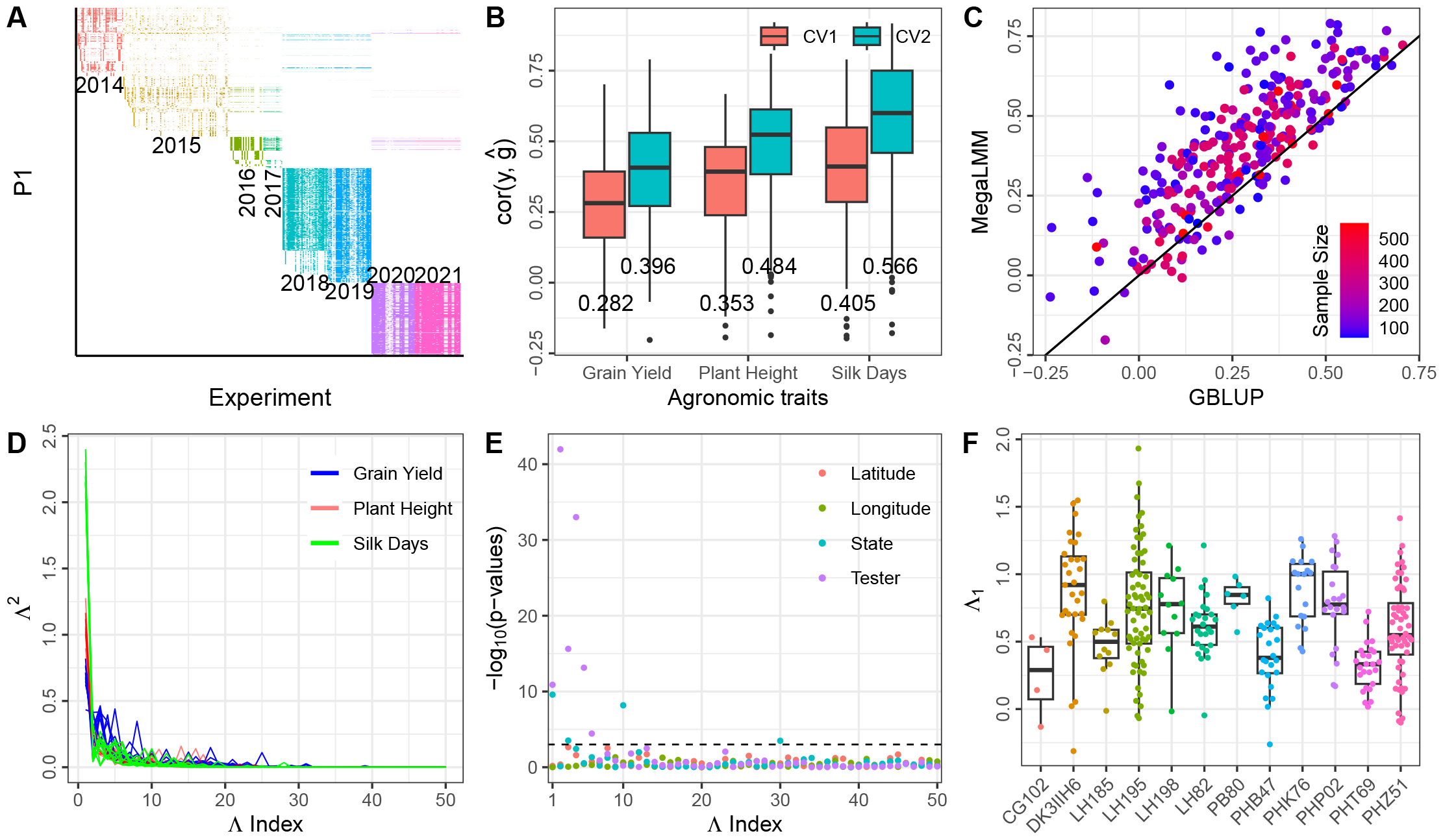
Data structure and predictive ability of the original MegaLMM model applied to the Genomes to Fields (G2F) maize hybrid dataset. (A) The data structure of the reshaped Genomes to Fields (G2F) phenotypic matrix, with inbred parent 1 (P1) in rows and experiments (combinations of location, year, and tester) in columns. Each cell in the matrix is filled with the least squares mean estimate of yield for a single hybrid genotype in a single experiment, with different colors indicating yield estimates from different years. (B) Boxplots of genomic prediction for CV1 and CV2 using the original MegaLMM model. Each point within a boxplot represents predictive ability for a specific experiment. The mean predictive ability for each trait within each scenario is shown below the corresponding boxplot. (C) Scatterplot of MegaLMM versus GBLUP predictive abilities for CV2 using the original MegaLMM model for Grain Yield. Each point represents a specific experiment. (D) Line plots showing magnitudes of squared factor loadings (**Λ**^2^). Each line represents the Λ^2^ per factor distribution in one MegaLMM chain for a specific agronomic trait. Different colors indicate distinct traits. We specified that MegaLMM should estimate 50 factors per dataset and ran 10 replicate MCMC chains per dataset. (E) Distribution of (*−*log_10_(*p−* values)) (y-axis) of regressions of factor loadings (x-axis) on latitude, longitude, state, and tester. (F) Boxplots of factor loadings for the first factor grouped by tester.

We used 5-fold cross-validation to measure the genomic predictive ability of the original MegaLMM model for each of the three agronomic traits separately (Silk Days, Plant Height, and Grain Yield) when trained on data from all 302 experiments and evaluated using the 20% of trait values withheld as validation data in each individual experiment. For sparse testing applications (predicting trait values for hybrids observed in some experiments but not others (CV2), estimated predictive abilities averaged *r* = 0.40 *−* 0.57 across the three traits based on a meta-analysis accounting for measurement error (Figure 2B), an average improvement of *r* = 0.12 *−* 0.19 across traits relative to predictions based on univariate GBLUP models trained on each experiment individually (Supplemental Figure S1). For trait value predictions of new hybrids with no observations in the training data (CV1), estimated average predictive abilities ranged from *r* = 0.28 *−* 0.41 for the three traits (Figure 2B), an average improvement of *r* = 0.01 *−* 0.03 over the univariate GBLUP models (Supplemental Figure S1). In almost every trial, estimated accuracies were higher for sparse testing applications (CV2) (Figure 2C). These results parallel our earlier results applied to the first four years of the G2F dataset (Runcie *et al*. 2021).

To investigate why MegaLMM improved over univariate approaches despite using the same genetic predictors, we extracted posterior means of the latent factor score 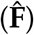 and factor loadings 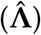 matrices. While these parameters are not always robustly identified in factor models like MegaLMM, the priors we use for elements of **Λ** tend to make specific factor orientations more reproducible. We allowed MegaLMM to learn 50 factors per dataset, but specified through our prior that the relative importance of factors should decrease rapidly across factor ranks. This effectively “turns off” many factors that are not needed when all loadings are shrunk close to zero. Applied to these three datasets, MegaLMM learned *∼* 13 *−* 20 factors per trait, each with at least one posterior mean importance weight (value of **Λ**) that explained *>* 1% of the trait variance. Distributions of weights across factors from 10 randomly chosen MegaLMM chains are shown in Figure 2D.

To explore whether candidate ECs such as the identity of the Tester or the geographic location, could serve as predictors of the importance weights for these factors, we regressed the posterior mean values of each row of **Λ** on latitude, longitude, state, or Tester. Among the 50 factors derived from the Grain Yield data in a single MegaLMM chain, four weight vectors were significantly associated with state and six were significantly associated with testers, based on Bonferroni-adjusted P-values *<* 0.05 (Figure 2E). Some factors displayed moderate correlations with either latitude or longitude; however, these correlations were not deemed statistically significant based on Bonferroni-adjusted P-values < 0.05 (Figure 2E-F). Therefore, factor loading weights are somewhat predictable based on known features of each trial and a hierarchical model including these ECs as features may be successful.

### Environmental Covariates enable MegaLMM to make accurate Genomic Prediction in new environments

We extended MegaLMM to additionally take as inputs ECs for any number of environmental features and use these as priors for factor loadings. We designed four Cross-Validation experiments to evaluate whether ECs could enable accurate genetic values predictions in new experiments in increasingly challenging prediction scenarios: 1) *NewTrial*: Can we predict genetic values in new trials, *e*.*g*. future trials that re-use previously observed testing locations and hybrids? 2) *NewState*: Can we predict genetic values in trials (of previously observed hybrids) grown in new geographic locations not near any existing trials *e*.*g*. in a new state? 3) *NewTester*: Can we predict genetic values of new hybrids (created with previously used P1s), *e*.*g*. new experiments in the same trials? 4) *NewGenoNewYear*: Can we predict the genetic values of previously unobserved hybrids (derived from neither same P1 nor same Tester) in new years. This scenario is motivated by the introduction of new inbred lines every two years, as depicted in Figure 2A.

Since in each case, the target experiments shared either geographic proximity (same state), or genetic similarity (same Tester) with trials in the training data, we first tested whether the extended MegaLMM model could improve genetic value predictions in these contexts using simple categorical ECs – the identity of the state and the identity of the Tester used in each experiment. We call this model S+T::S+T, signifying that the predictors S (State) and T (Tester) were used both in training (before the “::”) and as feature values for prediction (after the “::”).

As expected, genomic predictive abilities of MegaLMM declined across the four prediction scenarios for Grain Yield (Figure 3). Surprisingly, for Plant Height and Silk Days, the genomic predictive abilities of *NewTrial* were slightly lower than those of *NewState*, although the differences were not significant. Across the three agronomic traits, average genomic predictive abilities using the ECs ranged from 0.363 to 0.565 for *NewTrial*, 0.356 to 0.572 for *NewState*, and 0.221 to 0.438 for *NewTester* (Figure 3). The *NewGenoNewYear* scenario exhibited the lowest predictive ability, ranging from 0.080 to 0.264. We note that direct comparisons among the four scenarios are not entirely equitable due to subtle differences in the composition of training and testing sets associated with each scenario. Nevertheless, these comparisons offer an initial insight into the levels of predictive ability in each prediction scenarios.

**Figure 3.**
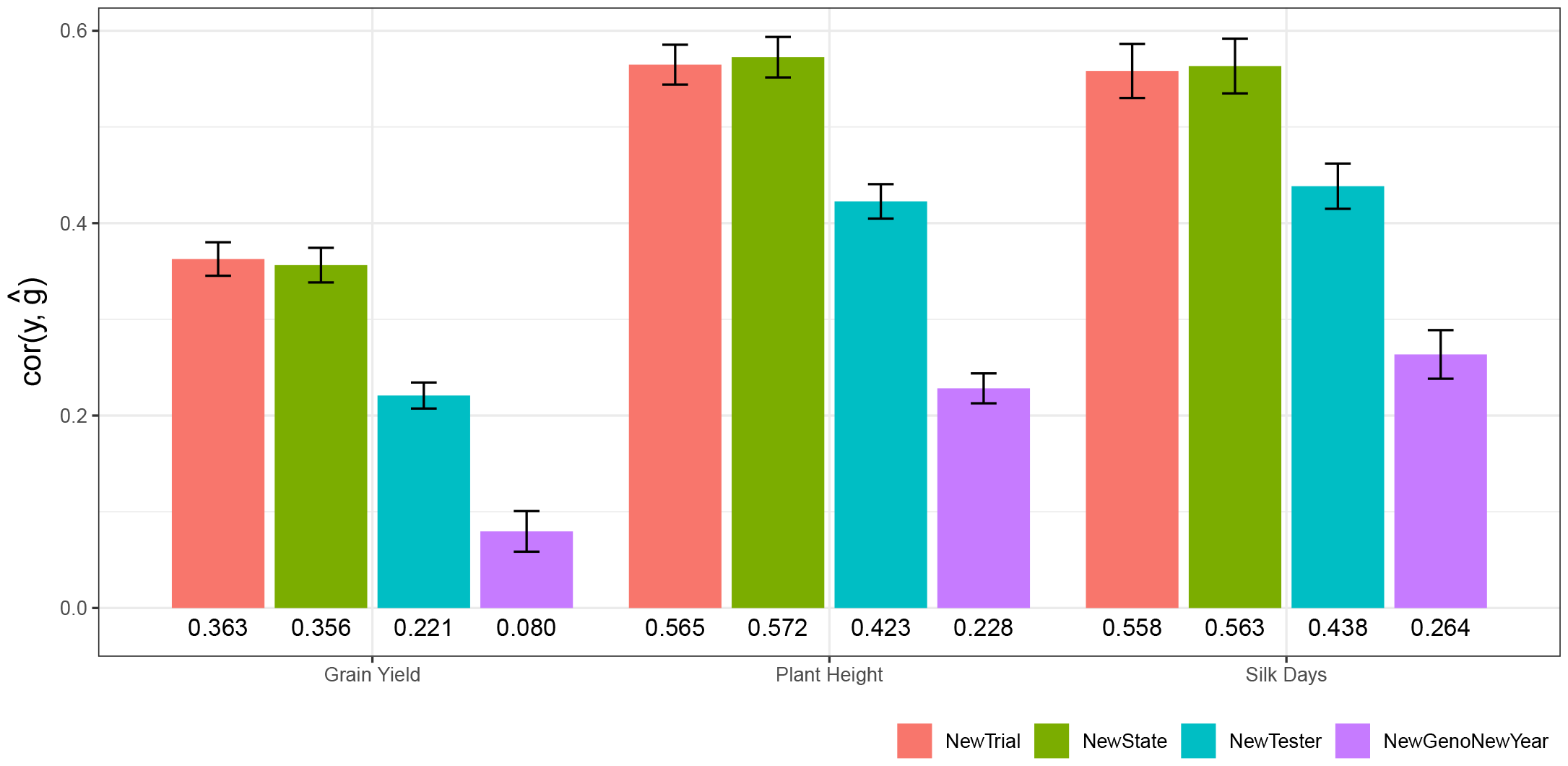
The extended MegaLMM model using ECs has moderate predictive abilities in new environments across four prediction scenarios for three agronomic traits (Silk Days, Plant Height, and Grain Yield). Each bar represents the estimated mean predictive ability of the extended MegaLMM model using State (“S”) and Tester (“T”) IDs as predictors (*i*.*e* “S+T::S+T” model) across individual experiments in a specific prediction scenario. The mean prediction accuracies for each trait within each scenario are shown below the corresponding barplot. Colors indicate prediction scenario. Error bars represent 95% confidence intervals of the mean, estimated by meta-analysis accounting for the size of each individual experiment. Note that EC “S” has no impact on factor loading predictions in the NewState scenario, and “T” has no impact on loading predictions in either NewTester or NewGenoNewYear scenarios.

To test if these predictive abilities were higher than could have been achieved using univariate GBLUP, we considered two univariate prediction strategies for the new environments: i) forming predictions of all candidate hybrids individually in each training experiment and then averaging predictions across all experiments into a single constant prediction to be applied to each new experiment, or ii) repeating this procedure but only for “similar” experiments, where we defined similarity as either experiments from the same state, or experiments using the same Tester. We ran both strategies and identified which produced more accurate predictions on average across all test experiments in a particular scenario. We used a similar procedure to select the most accurate extended MegaLMM model (i.e. experiment-specific or experiment-average predictions). We then compared the accuracies of the best MegaLMM model with the best GBLUP model for each experiment for each trait within each prediction scenario. Consistently, across all traits and nearly all scenarios, estimated mean prediction accuracies of MegaLMM were significantly higher than those of GBLUP (p-value < 0.01) (Figure 4). The exceptions were for the scenario of *NewGenoNewYear* for Grain Yield and Plant Height, where there was no significant difference between MegaLMM and GBLUP.

**Figure 4.**
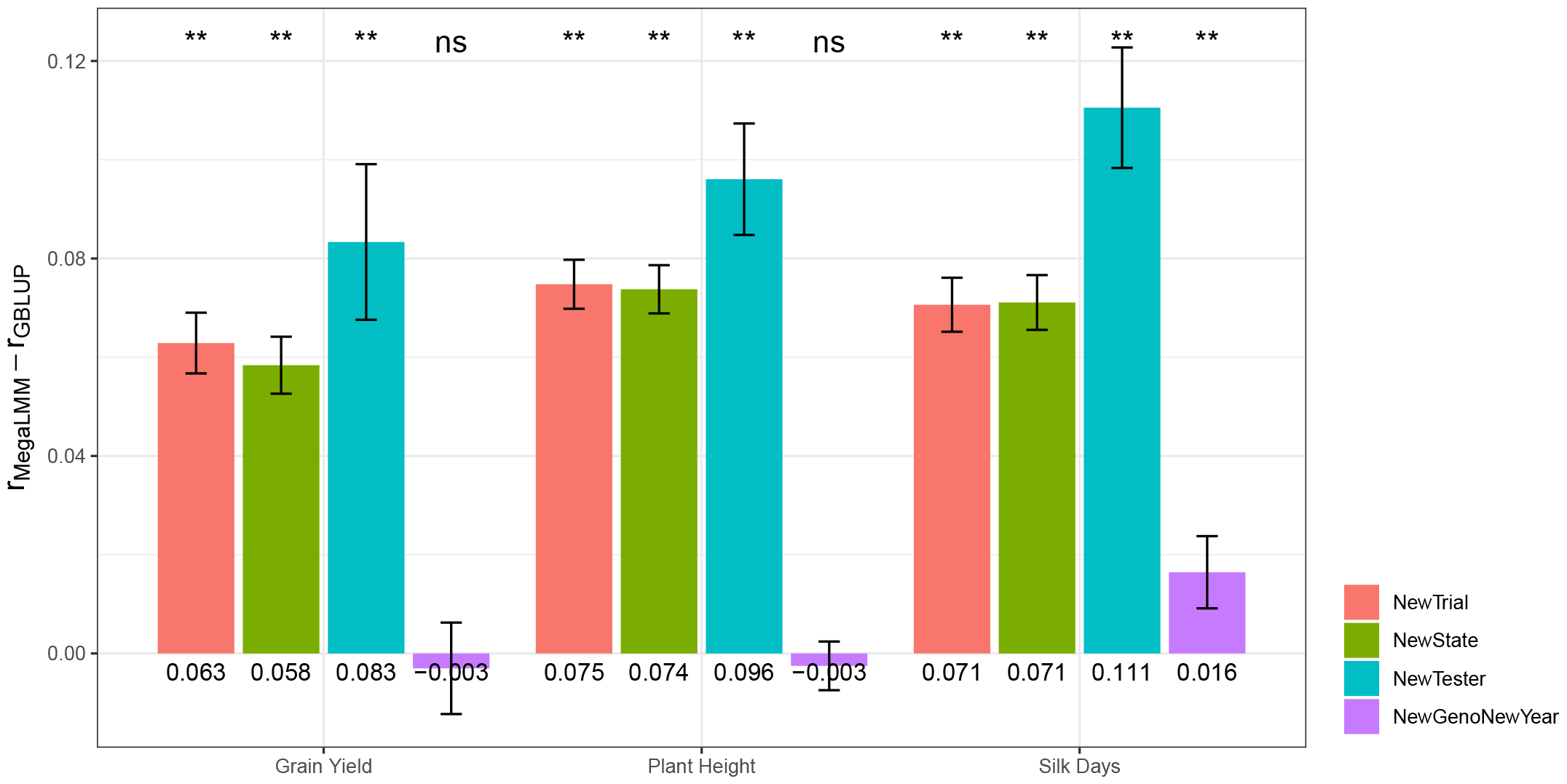
Predictive abilities of the extended MegaLMM model improve relative to univariate GBLUP prediction across most scenarios for three agronomic traits (Silk Days, Plant Height, and Grain Yield). Each bar represents the mean difference in predictive ability between MegaLMM and GBLUP in specific scenarios. Colors indicate prediction scenario. Error bars represent 95% confidence intervals of the difference in mean predictive ability between MegaLMM and GBLUP, estimated by meta-analysis accounting for the size of each individual experiment. Significance levels from a meta-analysis, are indicated above each barplot and mean differences in predictive ability for each trait within each scenarios are presented below the respective barplot.

These results confirm that MegaLMM’s predictions remain better than univariate predictions, even in new environments. However, it could be that this improvement is due to MegaLMM’s ability to empirically learn covariances among experiments, not the additional information provided by the ECs. In fact, even if we run MegaLMM without ECs and use the univariate strategy of simply averaging predictions across all training experiments, MegaLMM’s predictions in new experiments are considerably more accurate than the univariate ones (O::O vs GBLUP_O::O, Figure 5). To measure the additional benefit of the ECs, we ran prediction models using the ECs only as a prior but predicting based on the experiment-average as above (S+T::O), and using the ECs both as prior and for prediction (S+T::S+T). Note that in some scenarios, either the S or the T predictors were uninformative because the test values were not present in the training experiments, and so these predictors were dropped.

**Figure 5.**
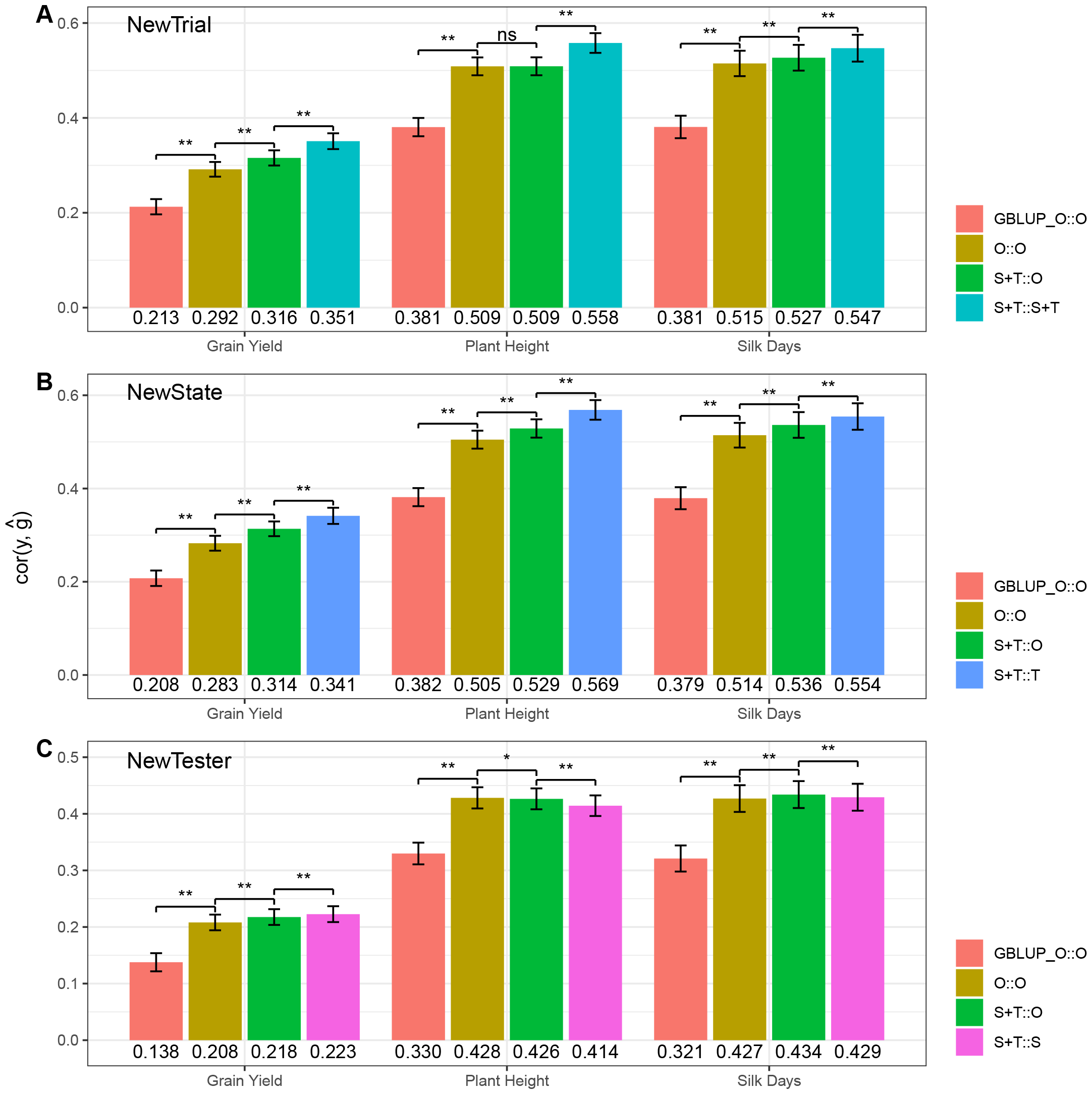
Environmental Covariates improve MegaLMM’s predictions in new environments relative to model that only use historical data, across three agronomic traits (Silk Days, Plant Height, and Grain Yield) and in three distinct prediction scenarios: (A) NewTrial, (B) NewState, and (C) NewTester. Each bar represents the estimated mean predictive ability of a specific model across individual trials. The mean prediction accuracies for each trait within each scenario are shown below the corresponding barplot. Colors indicate the prediction model. Error bars represent 95% confidence intervals of the mean, estimated by meta-analysis accounting for the size of each individual experiment. We show results from models with increasing complexity, starting with a univariate model, denoted GBLUP_O::O, based on GBLUP predictions obtained from averaging prediction made in individual experiments across all training experiments without using ECs, the original MegaLMM model that does not use ECs (O::O), a version of the extended MegaLMM model that uses ECs as priors but bases predictions on the average of predictions from each training trial without further using the ECs (S+T::O), and the full extended MegaLMM model that used ECs both as priors and as predictors (S+T::S+T). EC variables are denoted “S” for State, “T” for Tester, and “O” for empty ECs.

Across all three prediction scenarios, the S+T::O model significantly improved genomic predictive ability for all three agronomic traits, except for Plant Height in *NewTrial* and *NewTester*, where the S+T::O model’s accuracy was either identical or slightly lower than that of the O::O model (Figure 5A,C). These results suggest that the inclusion of S+T as a factor loading prior contributes to enhanced genomic prediction in new environments.

In the prediction scenarios of *NewTrial* and *NewState*, the S+T::T model significantly improved genomic predictive ability compared to the S+T::O model for all three traits (Figure 5A,B), indicating that incorporating the tester as a predictor enhances genomic predictive ability in new environments in cases where the same tester was used in training experiments. However, in the *NewTester* scenario, the S+T::S model outperformed the S+T::O model for Grain Yield but not for Plant Height and Silk Days (Figure 5C). These findings suggest that averaging similar experiments from the same Tester can enhance genomic predictive ability. However, when averaging similar experiments from the same State, the impact on genomic predictive ability varied, with sometimes showing slight improvements, but other times showing slightly decreased accuracy when predicting hybrid performance in new environments.

### Quantitative ECs can substitute of Qualitative ECs in Genomic Prediction for new environments

The above results demonstrate that MegaLMM can successfully use ECs to improve genomic predictive ability for new environments (Figure 5). However, these models used only categorical predictors (state or Tester labels), which can only be used to make predictions in environments that share the same levels of these labels. In contrast, quantitative ECs, like temperature or precipitation, could be used to make predictions in any geographic location.

To test whether MegaLMM can effectively use quantitative ECs, we repeated the above cross-validation experiments but substituted the categorical ECs with quantitative ones. We replaced the “S” ECs with scaled eigenvectors from a set of 278 weather variables (“W”), and the “T” ECs with the scaled eigenvectors of a genomic relationship matrix (“K”) of the Testers computed from the same genotypic data used for the P1s. Specifically, we compared a W+T::W+T model with the S+T::T model in the *NewState* scenario and a S+K::S+K model with S+T::S model in the *NewTester* scenario. Since “S” ECs cannot contribute to predictions in the *NewState* scenario but “W” can, and since “T” cannot contribute to predictions in the *NewTester* scenario but “K” can, we hypothesized that the use of quantitative ECs (“W” and “K”) would improve genomic prediction. However, only the S+K::S+K model significantly improved genomic predictive abilities compared to the S+T::S model in the *NewTester* scenario (Figure 6B). Other MegaLMM models with quantitative ECs showed either similar or slightly lower genomic predictive ability compared to their counterparts with qualitative ECs (Figure 6).

**Figure 6.**
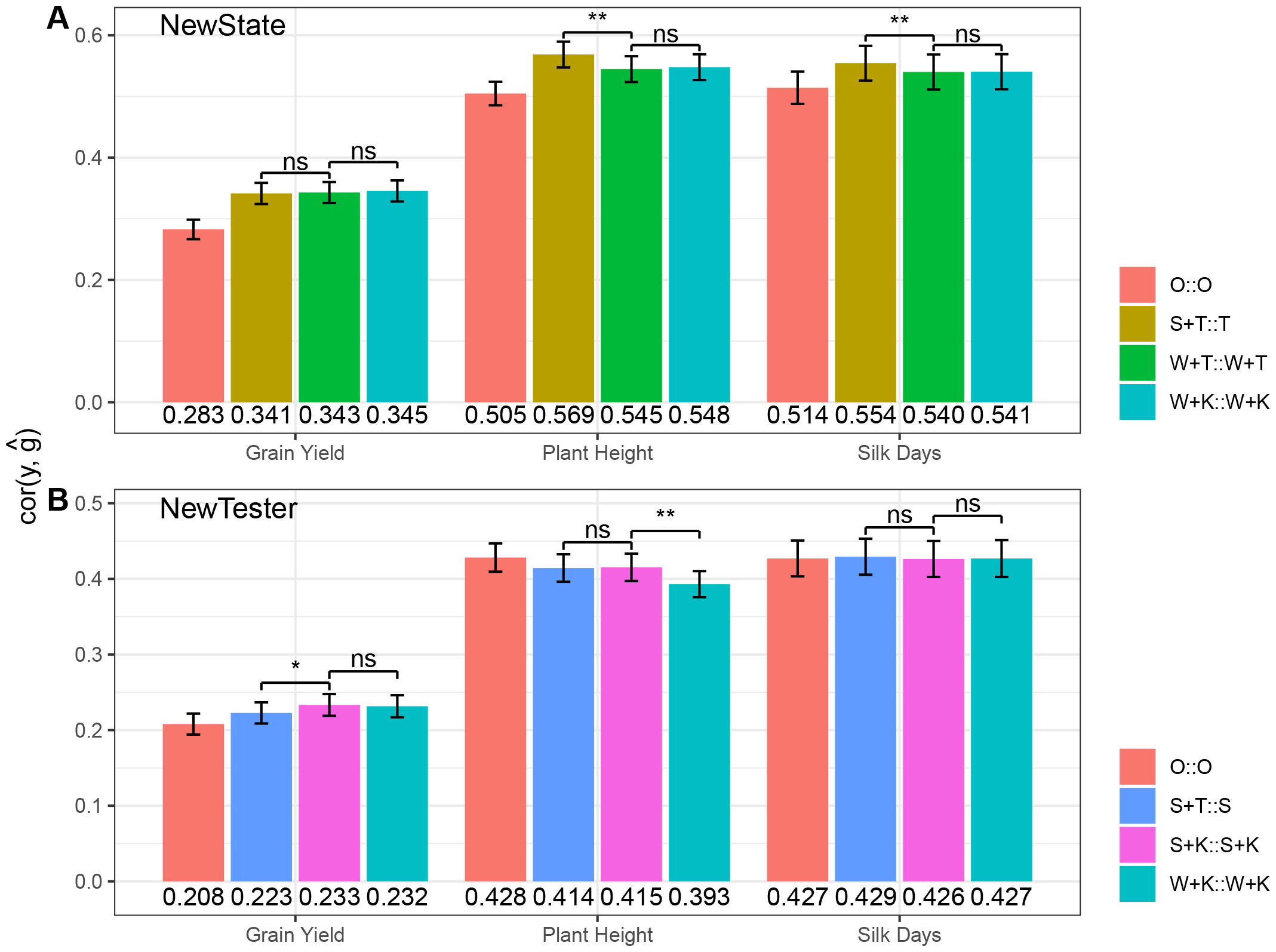
MegaLMM Model Comparison across three agronomic traits (Silk Days, Plant Height, and Grain Yield) using high-dimensional Environmental Covariates: (A) NewState and (B) NewTester. Each bar represents the estimated mean predictive ability of a specific MegaLMM model across individual trials. The mean prediction accuracies for each trait within each scenario are shown below the corresponding barplot. Colors indicate the prediction model. Error bars represent 95% confidence intervals of the mean, estimated by meta-analysis accounting for the size of each individual experiment. MegaLMM models are denoted by a combination of a factor loading prior and a factor loading predictor, separated by “::”, where “S” denotes State, “T” denotes Tester, “W” denotes Weather data, “K” denotes genomic relationship among Testers.

Subsequently, we replaced categorical ECs with quantitative ECs in MegaLMM models for the three traits in both scenarios. Specifically, we substituted “K” for “T” in the W+T::W+T model to obtain the W+K::W+K model in the *NewState* scenario and substituted “W” for “S” in the S+K::S+K model to get the W+K::W+K model in the *NewTester* scenario. We found that W+K::W+K model (with only quantitative ECs) performed just as well as W+T::W+T (with the “T” categorical EC) in the *NewState* scenario, and the W+K::W+K model (with only quantitative ECs) was only slightly inferior to the S+K::S+K model (with the “S” categorical EC) for Plant Height in the NewTester scenario (Figure 6). These results suggest that kinship and weather data can effectively substitute for categorical labels when the states or testers have been observed in the training data. This shows that MegaLMM can effectively use quantitative ECs, but suggests that the quality of the quantitative ECs we used in our analysis may have been too low to be useful in this analysis.

## DISCUSSION

### Insights into the use of Environmental Covariates for genomic prediction for new environments

We developed an extended MegaLMM model with EC-based priors to predict genetic values in new environments. MegaLMM is based on a factor-analytic model, and allows users to model a large number of latent factors underlying variation in genetic values in each environment. The ECs help the model learn the importance weights for each factor in each environment, and provide coefficients necessary to predict importance weights - and therefore genetic values - in new environments. Overall, we found the ECs significantly improved MegaLMM predictive ability in most scenarios relative to the performance of the MegaLMM base model (O::O, Figure 5). However, compared to the improvement of the MegaLMM base model over a univariate GBLUP approach (Figure 4), the improvement due to the incorporation of ECs was less dramatic. These results highlight several important points.

First, even though neither the MegaLMM base model nor univariate GBLUP models can directly produce predictions of genetic values in new environments, we found that a simple post-processing of their predictions across the MET experiments (*i*.*e*. old environments) could result in reasonably accurate genetic value predictions on average in new environments. Specifically, the average predicted genetic values across the MET experiments were correlated with observed values in most new environments. In some cases, but not always, we could improve predictions by clustering the MET experiments either by geography (State) or Tester and only averaging the genetic value estimates within a cluster when predicting genetic values in new experiments in the same cluster. This latter approach can be thought of as a non-parametric approach for using the ECs, and is equivalent to factorial regression (Denis 1988; Piepho *et al*. 1998) approaches using categorical ECs as dummy variables. One reason that such constant (*i*.*e*. not environment-specific) predictions can be successful in this dataset is that trait values are positively correlated between most environments (Supplemental Figure S2), diminishing the potential benefit of forming unique predictions in each new environment. Thus, while models do detect significant G*×*E in this dataset (Rogers *et al*. 2021; Lopez-Cruz *et al*. 2023), the magnitude of the G*×*E variance is not large relative to genetic main effect variance. G*×*E models necessarily have larger prediction variances because they try to make more specific predictions, and unless the actual G*×*E variance is large enough to counteract the reduced precision, “main effect” models will be more accurate (Weine *et al*. 2023). One possible reason for the relatively low importance of G*×*E in this dataset is the wide diversity among hybrids, including some relatively low-performing hybrids with poor trait values in most environments. If only elite hybrids had been used, the relative importance of G*×*E prediction might have been higher.

Second, ECs are useful for learning the model’s parameters even if not used for prediction. We found that when we used ECs as priors during model training, the correlation between the averages of predicted genetic values across the MET experiments and the observed phenotypes in new environments was typically higher than with the MegaLMM base model which did not use the ECs (Figure 5). In this case, we did not use the ECs to make predictions tailored to each new environment, yet still found the ECs useful. Here, the ECs may help the model learn correlations between pairs of trials that do not share many hybrids in common, so there is little data to learn correlations empirically, but which do share values of ECs. This suggests that ECs may be especially useful when METs are very sparse and perhaps even unconnected – containing trials without *any* overlapping hybrids. This improvement was not apparent within the MET experiments themselves (in terms of accuracy measured by CV2), probably because the residual genetic terms (*U*_*R*_) were able to make sufficiently accurate predictions.

Third, successfully predicting genetic values in new environments may require both higher-quality ECs and many more MET experiments. While this data set is large, composing 302 experiments, it contains only 195 trials in different site-years to learn regressions on ECs like weather, only 37 locations to learn regressions on ECs like geography, climate, and soil, and only 12 testers to learn regressions on genetic markers of each tester. Genomic prediction models (in a single environment) generally require hundreds of genotypes to effectively learn allele-phenotype correlations (Jannink *et al*. 2010) because genotypes are the unit of replication of alleles in these models. Since experiments are replicates of environmental variables in G*×*E models, and because the environmental drivers of performance are likely similarly complex to genetic drivers, hundreds of experiments are probably needed to adequately model G*×*E in new environments. Nevertheless, we showed that MegaLMM could successfully use high-dimensional ECs (from weather or tester genotypes) to make accurate predictions, at least when the new environments were closely related to existing environments (same states or same testers). However, more informative ECs, such as ECs derived from crop growth models (Heslot *et al*. 2014; Rincent *et al*. 2019) may help reduce the dimensionality burden, making G*×*E modeling more efficient.

### Comparison with other approaches for predicting genotype-environment interactions

Compared with previous statistical models that use ECs for predictions in new environments, the extended MegaLMM model offers several statistical and practical advantages, including the ability to use high-dimensional ECs, regularization through a moderate number of latent factors, and the ability to fit phenotypic data from very large and very sparse METs.

The ability to simultaneously use high-dimensional ECs for prediction should be useful when multiple environmental variables simultaneously impact the variation in genetic values across a TPE. In the extended MegaLMM model, we use regularized regression to provide robust inference across high-dimensional ECs. This contrasts with the CERIS-JGRA method of Li *et al*. (2021) which searches among candidate ECs for a single EC that is best, and then bases predictions on this single EC. Also, while the CERIS-JGRA method selects an EC based on the ability to predict phenotype means across trials, the extended MegaLMM prioritizes ECs based on their usefulness for distinguishing patterns of covariance among trials, which is more directly applicable to breeding.

The ability to robustly use high-dimensional ECs is not unique to the extended MegaLMM model. The GBLUP-based reaction norm model, as demonstrated by Jarquín *et al*. (2014), can also use high-dimensional ECs, using kernel functions to turn the EC matrices into distance matrices. Costa-Neto *et al*. (2021) also uses kernel methods for model G*×*E from METs. A limitation of this approach is that training the kernel functions themselves is computationally expensive, so these methods use fixed kernel functions which prevents learning weights among the ECs. Tuning parameters of the kernel functions is possible in these methods, but the same tuned kernels would apply to all trials. In contrast, the extended MegaLMM model can learn different EC weights for each latent factor, providing an additional level of flexibility and opportunity for statistical learning of G*×*E patterns.

Much of MegaLMM’s statistical and computational efficiency comes from its latent factor model architecture. Many other models also use factor-analytic models for G*×*E prediction. For example, the AMMI model is a factor model (but with fixed factors, Rincent *et al*. (2019)), and Cullis *et al*. (2014) and Heslot *et al*. (2014) also proposed factor-analytic models for METs. The advantage of factor-analytic models is that they model correlated traits with a small number of parameters relative to the number of covariances among pairs of trials, providing statistical robustness, and remove the need to invert large covariance matrices, alleviating computational limitations. However, MegaLMM is unique in its ability to fit relatively large numbers of latent factors. Most prior applications of factor-analytic models have handled only 1-3 factors and can fail to converge if run with more factors. For example, Schulz-Streeck *et al*. (2013) found that the factor analytic structure failed to converge when fitting a marker-by-environment interaction model, and Rogers *et al*. (2021) found that models with more than one FA factor for environments, in combination with either additive or dominance relationships, failed to converge when fitting a subset of G2F data. Our analysis of the maize G2F dataset used 50 factors and found that 13-20 factors significantly contributed to trait performance prediction across experiments for three agronomic traits (Figure 2D). This suggests that more factors can be beneficial for accounting for varying sources of G*×*E variation in large METs.

Finally, our case study was a MET with 302 experiments and *∼*87% missing values, yet MegaLMM was able to return predictions in *∼*3 hours (with 20 CPU cores). The ability to fit such sparse data is an advantage over the AMMI models and other matrix-based (i.e. **Y** is treated as a matrix) models that require complete data. Also, the efficiency in fitting data with large numbers of traits makes MegaLMM flexible for modeling complex experimental features like management characteristics which can be important contributors to G*×*E (Cooper *et al*. 2021). Modeling management in addition to environmental drivers complicates reaction norm models because of the need to specify many interaction terms, making models unwieldy. In contrast, as a correlated traits model, MegaLMM does not explicitly require an interaction term to model G*×*E*×*M effects. Integrating management characteristics into the MegaLMM model involves simply expanding the columns in the multivariate response matrix, with columns representing combinations of environmental types and management. Mathematically, the process of solving the linear mixed model equations and estimating parameters remains unchanged.

### Insights into modeling G×E in the maize hybrid breeding system

Contrasting with most other analyses of the G2F maize hybrid dataset (Rogers *et al*. 2021; Lopez-Cruz *et al*. 2023), we divided each trial into multiple separate experiments based on the identity of different testers used to create each hybrid, and then modeled the covariances among these experiments. There are both practical and statistical benefits to doing this. On the practical side, focusing on within-tester-family predictive ability aligns our approach with maize hybrid breeding strategies. In maize hybrid breeding, germplasm is organized into two major heterotic pools, and inbred lines are developed within these pools. The newly created inbred lines are initially evaluated and selected by crossing them with suitable testers from complementary heterotic pools. Subsequently, they are further crossed with a larger number of newly created lines from the opposite heterotic pool for evaluation for potential commercial use (Cooper *et al*. 2014). In our analysis, we placed inbred lines, rather than hybrids, as rows of our data matrix **Y**, with columns representing combinations of Tester and environment. Thus, our genomic predictions are best considered genetic values of inbreds conditional on specific Testers and environments.

On the statistical side, modeling the covariance among hybrids from different testers allows modeling of Tester-inbred genotype interactions, and therefore produces more accurate within-tester-family predictions when these interactions are important. Since MegaLMM scales very efficiently with numbers of experiments, there is little downside to breaking trials into multiple experiments, particularly when we can include prior information through ECs to partially pool information across experiments when they are closely related. We found that Tester identity was the most useful EC among experiments (comparing results from the *NewState* scenario where Tester ID could be used for prediction in new environments, to results from the *NewTest* scenario where Tester ID was not available for prediction, Figure 3), suggesting that the ranking of inbreds did change considerably when crossed to different Tester. However, this result should be interpreted with caution because the importance of Tester ID in this dataset is partially confounded with both geographic structure among trials and population structure within the populations of inbreds (P1s), as discussed by Lopez-Cruz *et al*. (2023).

In summary, we present an extended version of MegaLMM that can predict the genetic architecture of new traits based on trait-specific prior data. This is a significant advancement of the MegaLMM method, opening the possibility of many types of novel applications. We focus here on the application of modeling genotype-environment interactions in multi-environmental trials in plant breeding, where we consider each trial a new trait, and use environmental data as prior predictors of the patterns of genotype-environment interactions. We expect that many other applications of this extended MegaLMM model are possible both in plant breeding and in other fields where large linear mixed models can be applied.

## METHODS

### Original MegaLMM Model

The original MegaLMM “correlated-traits” model of a MET is specified as:

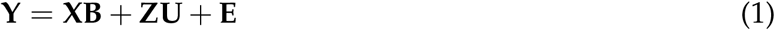

where:

**Y** is an *n × t* phenotypic matrix for a trait of interest measured on *n* experimental genotypes grown in *t* trials, potentially with a large percentage of missing values,

**X** is an *n × p* incidence matrix for fixed effects such as an intercept,

**B** is a corresponding *p × t* matrix of fixed effects for each trial,

**Z** is an *n × q* incidence matrix for random effects, in this case the identities of each inbred parent,

**U** is a corresponding *q × t* matrix of random effects, in this case additive genetic values, for each trial,

**E** is an *n × t* matrix of residuals for each genotype in each trial.

Fitting Eq. (1) is challenging because the columns of **U** and **E** are correlated. To address this issue, Runcie *et al*. (2021) developed a new statistical framework, MegaLMM, based on a factor analytic model, which decomposes the correlated traits model into a two-level hierarchical model.

In level 1, the phenotypic matrix **Y** is decomposed into two components:

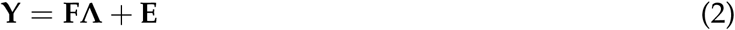

where:

**F** is an *n × k* latent factor matrix,

**Λ** is a *k × t* loading matrix,

**E** is an *n × t* residual matrix of residuals for each trial.

Intuitively, *k* latent factors can be interpreted as *k* unobserved traits across each individual that are constant across experiments, and the factor loadings represent the relative importances of each of these *k* unobserved traits on the focal trait value in each experiments.

In level 2, each of the *k* latent factors in the **F** matrix and each of the *t* residual traits in the **E** matrix are independently fitted with standard univariate linear mixed models:

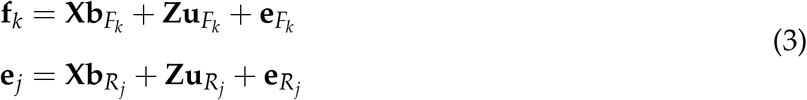

where:

**f**_**k**_ and **e**_*j*_ are *n ×* 1 vectors for the *k*th latent factor trait and the *j*th residual trait, respectively.

**X** is an *n × p* incidence matrix for fixed effects,

**Z** is an *n × q* incidence matrix for random effects,

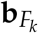 and 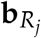 are *p ×* 1 vectors of fixed effects for the *k*th factor and *j*th residual trait, respectively,

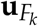 and 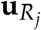 are *n ×* 1 vectors of random effects for the *k*th factor and *j*th residual trait, respectively,

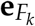 and 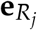 are *n ×* 1 vectors for residuals.

The distributions of random effects are specified as:

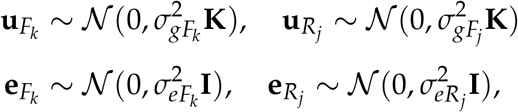

where:

**K** is the pairwise genomic relationship matrix between old genotypes that is estimated with genetic molecular markers,

**I** is the identity matrix,

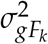 is the genetic variance components associated with the kth latent factor,

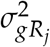 is the genetic variance components associated with the jth residual trait,

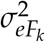 is the residual variance components associated with the kth latent factor, and

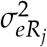 is the residual variance components associated with the jth residual trait.

All parameters of MegaLMM are estimated using a Gibbs sampler as described in (Runcie *et al*. 2021).

### Extensions to Predict Trait Performance in New Environments

The original MegaLMM model lacked the capability for making predictions in new environments because elements of the environment-specific weights matrix **Λ** were independent in the prior and thus could only be learned based on correlations between records in different observed environments. Our extended MegaLMM model replaces the original prior on **Λ** with a prior of the following form:

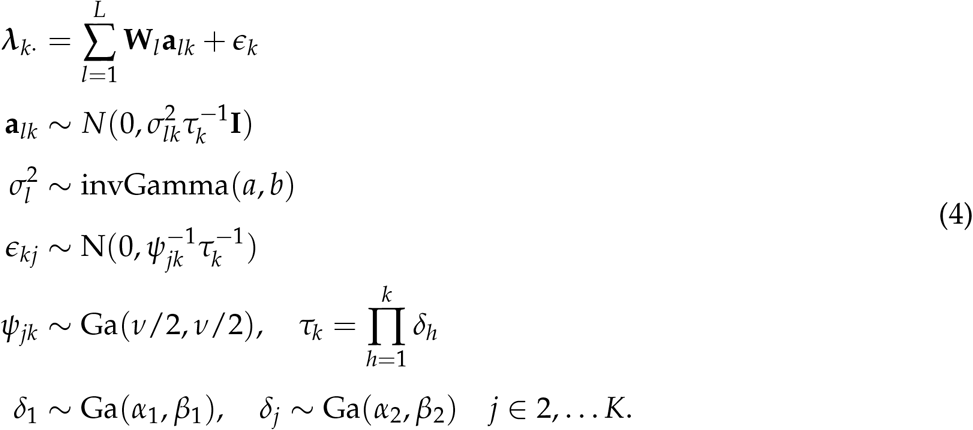

where ***λ***_*k·*_ is a row of **Λ** representing the relative importance weights of latent factor *k* across environments. We model this vector as a regression on ECs, represented as *L* design matrices **W**_*l*_, *l ∈* 1 … *L*, for example **W**_1_ is usually a single column of 1’s representing an intercept, and in the MegaLMM_S+T model, **W**_2_ would be an incidence matrix of state identities, and **W**_3_ would be an incidence matrix of Tester identities. The regression coefficients are assigned independent normal priors with a variance that shrinks for higher order factors based on the precision parameter 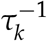. The residuals of this regression are assigned heavy-tailed t-distributed priors as in our earlier BSFG model (Runcie and Mukherjee 2013), which maintains the shrinkage of higher order factors towards zero. Parameters of this model for **Λ** are learned using the same Gibbs sampler steps as in the BSFG model (Runcie and Mukherjee 2013).

Using posterior samples of the regression coefficients 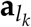, posterior predictions of genetic values in new environments can be formed as:

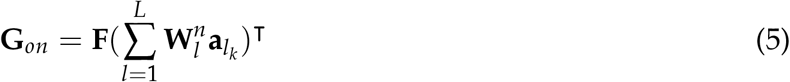

where:

**G**_*on*_ are posterior samples of the genetic value for old genotypes in new environments,

**F** are posterior samples of the latent factor matrix estimated from old grown in old environments,

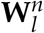 are values for ECs in the **W**_*l*_ matrix measured in new environments.

To form posterior predictions of genetic values for new genotypes, **F** in 5 is replaced with **F**_*n*_ = **K**_*no*_**K**^*−*1^**F**, where **K**_*no*_ is the is the pairwise genomic relationship matrix between new and old genotypes.

### Cross-validation scenarios for predicting experimental genotypes in old and new environments

#### CV1

We randomly divided experimental genotypes into five equal-sized folds within each environment. The partition of genotypes was consistent across all environments. During cross-validation, four folds were used for model training, and the fifth fold served as the validation set. This process was repeated five times until each of the five folds in each environment was used as the validation set.

#### CV2

Within each environment, we used the same genotype partition as CV1. However, we randomized the order of the five folds independently across environments. During cross-validation, four folds were used for training, and the fifth fold was used for validation. This procedure was repeated five times until each of the five folds within each environment served as the validation set.

#### NewTrial

Building on the CV2 training sets, we randomly divided all trials (i.e., location-year combinations) into five folds. Four folds were used for training, and the fifth fold was used for cross-validation. For each of the five distinct CV2 training sets, this process was repeated five times until each of the five folds of trials had been used as a validation set.

#### NewState

Following the CV2 training sets, we split all experiments by their respective States. We selected States with at least 9 experiments as testing sets, resulting in 14, 13, and 12 testing sets for Grain Yield, Plant Height, and Silk Days, respectively, for the G2F dataset. For each State in the testing set, all other States were used for training. For each of the five distinct CV2 training sets, this process was repeated 14, 13, and 12 times for Grain Yield, Plant Height, and Silk Days, respectively, until each set of testing experiments had been used as a validation set.

#### NewTester

Based on the CV2 training sets, we split all experiments by their testers, resulting in a total of 12 sets of testing experiments for the G2F data. Each set of testing experiments served as a testing set, and the remaining experiments were used for model training. For each of the five distinct CV2 training sets, this process was repeated 12 times until each set of testing experiments had been used for validation.

#### NewGenoNewYear

Using each of the CV2 training sets, we divided all experiments into four folds based on two-year intervals (2014-2015, 2016-2017, 2018-2019, and 2020-2021). Since hybrid compositions changed dramatically every two years, each fold contained almost entirely different sets of hybrids. To ensure no overlap between training and testing sets, we further excluded common hybrids from the testing set. Thus, each fold represented new genotypes tested in new environments. For each of the five distinct CV2 training sets, this process was repeated four times until each of the four folds of experiments had been used as a validation set.

### Estimating genomic prediction accuracies, their means and standard deviations

Within each experiment, predictive ability was estimated using the following equation:

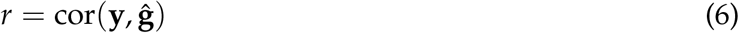

where:

**y** is a vector of adjusted phenotypic values, and

**ĝ** is a vector of predicted genotypic values.

For CV1 and CV2, within each experiment, we defined predictive ability as the mean correlation obtained from five validation sets. Similarly, for prediction scenarios of *NewTrial, NewState, NewTester* and *NewGenoNewYear*, within each experiment, we defined predictive ability as the mean of prediction accuracies obtained from five distinct validation sets, which originated from five distinct CV2 training sets.

Within each prediction scenario we estimated means and standard deviations of prediction accuracies over all experiments using a meta-analysis to different sample size with the Hunter and Schmidt-type approach (Schmidt and Hunter 2014) using the *escalc* and *rma* functions of the *metafor* R package (Viechtbauer 2010). This implements a random-effect meta-analysis with estimated standard errors of each individual correlation based on its own sample size. To test if one method produces higher correlations on average than another, we compared the two vectors of correlations using the *r*.*test* function of the psych R package (Revelle 2023), and extracted the estimated difference between the two methods for each trial as well as the standard error of this difference. We then used the *rma* function of the *metafor* package to compute a random effects meta-analysis of these differences weighted by the sample size of each trial. Finally, we estimated 95% confidence intervals (CI) of mean predictive ability within each prediction scenario with the following equation:

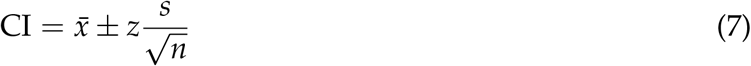

Where:

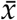 is the mean predictive ability,

*z* is the Z-score corresponding to the desired confidence level (for a 95% confidence level, z= 1.96),

*s* is the standard deviation of the prediction accuracies across experiments,

*n* is the total number of experiments within a prediction scenario.

### Phenotypic and genotypic analyses of G2F maize hybrid dataset

#### Plant Materials

The Genomes to Fields Initiative (G2F) is a multi-institutional, collaborative initiative to catalyze and coordinate research linking genomics and phenomics in maize to achieve advances that generate societal and environmental benefits (AlKhalifah *et al*. 2018). Since 2014, this project has evaluated approximately 180,000 field plots involving more than 5,000 corn hybrid varieties across more than 200 unique environments in North America. Our analyses used the G2F maize hybrid data collected between 2014 and 2021 and focused on three representative agronomic traits: Grain Yield: Measured in Mg per ha at 15.5% grain moisture (unit: Mg/ha), utilizing plot area without an alley; Plant Height: Quantified as the distance from the base of the plant to the ligule of the flag leaf, expressed in centimeters; Silk Days: Defined as the number of days elapsed after planting when 50% of the plants within a plot displayed visible silks.

#### Phenotypic Data Analysis

The initial 2014-2021 G2F phenotypic dataset comprises 217 unique trials with diverse field experiment designs. As more than 71.4% of the G2F data points were linked to 12 major hybrid testers (Lopez-Cruz *et al*. 2023), our analysis concentrated on these key tester families. Consequently, within each trial (i.e., a location::year combination), we split the trait data by Tester and refer to each partition as an experiment. We selected experiments composing a minimum of 50 hybrid genotypes for further analysis. Therefore, in our analysis we consider the Tester as a component of an environment.

Our pre-processesing of the raw phenotypic data from each trial included the following steps. First, we excluded tester families with fewer than 50 hybrid genotypes. Subsequently, we employed a two-step procedure to filter outliers. Initially, within each individual trial, outlier data points were eliminated based on the joint distribution of observed trait values across trials. Data points with an expected occurrence of less than 1, assuming a normal distribution, were flagged as outliers. Subsequently, outlier trials were identified based on the distribution of mean trait values across all trials. Trials with a population mean expected to occur less than 1 time, given a normal distribution, were classified as outliers. Following outlier removal, we retained 302, 278, and 231 experiments (i.e., tester families) for Grain Yield, Plant Height, and Silk Days, respectively, for downstream analysis.

To account for field design factors and obtain the best linear unbiased estimation (BLUE) of each hybrid genotype, we employed linear or linear mixed models, depending on available experimental design factors within each experiment. Experiments were categorized into four groups, each fitted with a different model:

- For experiments with >=2 replicates and >=2 blocks each, we used a linear mixed model: **y** *∼* Hybrid + Replicate + (1|Replicate:Block), where **y** represents observed phenotypic values, Hybrid and Replicate are fixed effects of hybrid genotypes and replicates, respectively, and (1|Replicate:Block) is the random effect of block nested within replicate.
- For experiments with >=2 replicates and only one block in each replicate, we employed a linear model: *y ∼* Replicate + Hybrid.
- In cases with only one replicate but multiple blocks in the replicate, we used a linear mixed model: *y ∼* Hybrid + (1|Block), where (1|Block) represents the random effect of block.
- For a few experiments with only one replicate and one block in the replicate, a linear model *y ∼* Hybrid was applied.

Linear mixed models were fitted using the *lmer* function in the R library *lme4* (Bates *et al*. 2015). Linear models were fitted with the *lm* function in the base R library (R Core Team 2023). The *predict* function from the base R library was employed to extract marginal BLUEs for each hybrid genotype in each environment.

Finally, we re-shaped all BLUEs for each hybrid genotype in each environment into a matrix with rows corresponding to each inbred Parent 1’s of the hybrid, and columns corresponding to the experiment IDs (i.e. location-year-tester combinations).

#### Genotypic Data Analysis

We received G2F genotypic data from the committee of The Genomes to Fields 2022 Maize Genotype by Environment Prediction Competition (Lima *et al*. 2023), who only provided genotypic data of hybrid genotypes. The 2014-2021 G2F inbred lines (Hybrid Parent 1s and testers) were sequenced with different technologies. The Maize Practical Haplotype Graph (PHG) database 2.1 was used for variant calling, which generated a genotypic dataset with 4,928 unique hybrid genotypes and 437,214 SNP sites. We first filtered the SNPs using the following criteria: (i) minor allele frequency (MAF) > 5%; (ii) maximum site missing rate < 20%, resulting in a dataset with 4928 unique hybrid genotypes and 324,323 SNP sites. We used a custom script to infer the P1 and Tester genotypes of each hybrid. Briefly, for each SNP in each hybrid, if the genotype was 0 or 2, we assigned this value to both parents. If the genotype was 1, either the P1 or the Tester must have the 1 allele. To decide, we compared the same locus to all other hybrids from the same tester. If any other hybrid had a 0 genotype at this locus, the Tester’s genotype must be 0, otherwise its genotype must be 1. For this analysis, we filtered out any hybrids where the tester was not replicated in at least one other hybrid.

Using the separate SNP genotype matrices of the P1s and the Testers, we computed additive genomic relationship matrices for each following VanRaden’s equation (VanRaden 2008) using the dogrm software package (Bellot *et al*. 2018).

#### Weather Data Analysis

The original weather environmental variable record was captured on a daily basis. Given the high correlation among these daily environmental variables, we conducted the following analyses to address redundancy in environmental covariates: (i) We computed the Daily Corn Growing Degree Days (GDD) between the planting and harvest dates for each trial using the formula: Daily Corn GDD (°F) = (Daily Maximum Temperature °F + Daily Minimum temperature °F) - 50 °F. If the daily maximum and/or minimum temperature was less than 50 °F (10 °C), it was adjusted to 50 °F. Similarly, if the daily maximum temperature exceeded 86 °F, it was capped at 86 °F. (ii) We computed the Accumulated Growing Degree Days (AGDD) and determined maize growth stages for each trial based on methodologies described by Widhalm (2014) and Nielsen (2019). This analysis identified 23 stages of maize growth, including 20 vegetative growth phases from emergence (VE), V1-V18, up to tassel formation (VT). For the reproductive phase, we consolidated R1, R2, and merged R3 to R6 into a single growth stage. (iii) We averaged 11 weather environmental variables (Supplemental Table S1) and GDD within the duration of each of the 23 growth stages. Moreover, AGDD and Accumulated Precipitation (APRE) of each trial were included as environmental covariates, recognizing temperature stress and water deficit as the two most important factors limiting crop growth and yield (Langridge *et al*. 2021). Ultimately, this process yielded 278 ECs.

## DATA AVAILABILITY

We obtained the G2F dataset from the committee of The Genomes to Fields 2022 Maize Genotype by Environment Prediction Competition, accessible on CyVerse under https://doi.org/10.25739/tq5e-ak26. The scripts used in this study are documented in the following GitHub repository: https://github.com/hh622/MegaLMM_New_Environments_Prediction_GenomesToFields. Additionally, the R package for extended MegaLMM can be found here: https://github.com/deruncie/MegaLMM/tree/restructure and will be moved to the ‘master’ branch and archived at Zenodo at time of publication.

## FUNDING

This research was funded by the National Institute of Food and Agriculture (NIFA)’s Agriculture and Food Research Initiative (AFRI) competitive grant, grant number 2020-67013-30904.

## AUTHOR CONTRIBUTIONS

D.E.R. and H.H. conceived the research and analyzed the data; H.H., D.E.R. and R.R. wrote the manuscript.

## ACKNOWLEDGMENTS

The authors acknowledge the committee of The Genomes to Fields 2022 Maize Genotype by Environment Prediction Competition for providing the maize hybrid datasets. Additionally, the authors thank Alencar Xavier from Corteva Agrisciences for his valuable advice on data analysis and manuscript editing.

## DECLARATION OF INTERESTS

No conflict of interest declared.

## SUPPLEMENTAL INFORMATION

**Figure S1.**
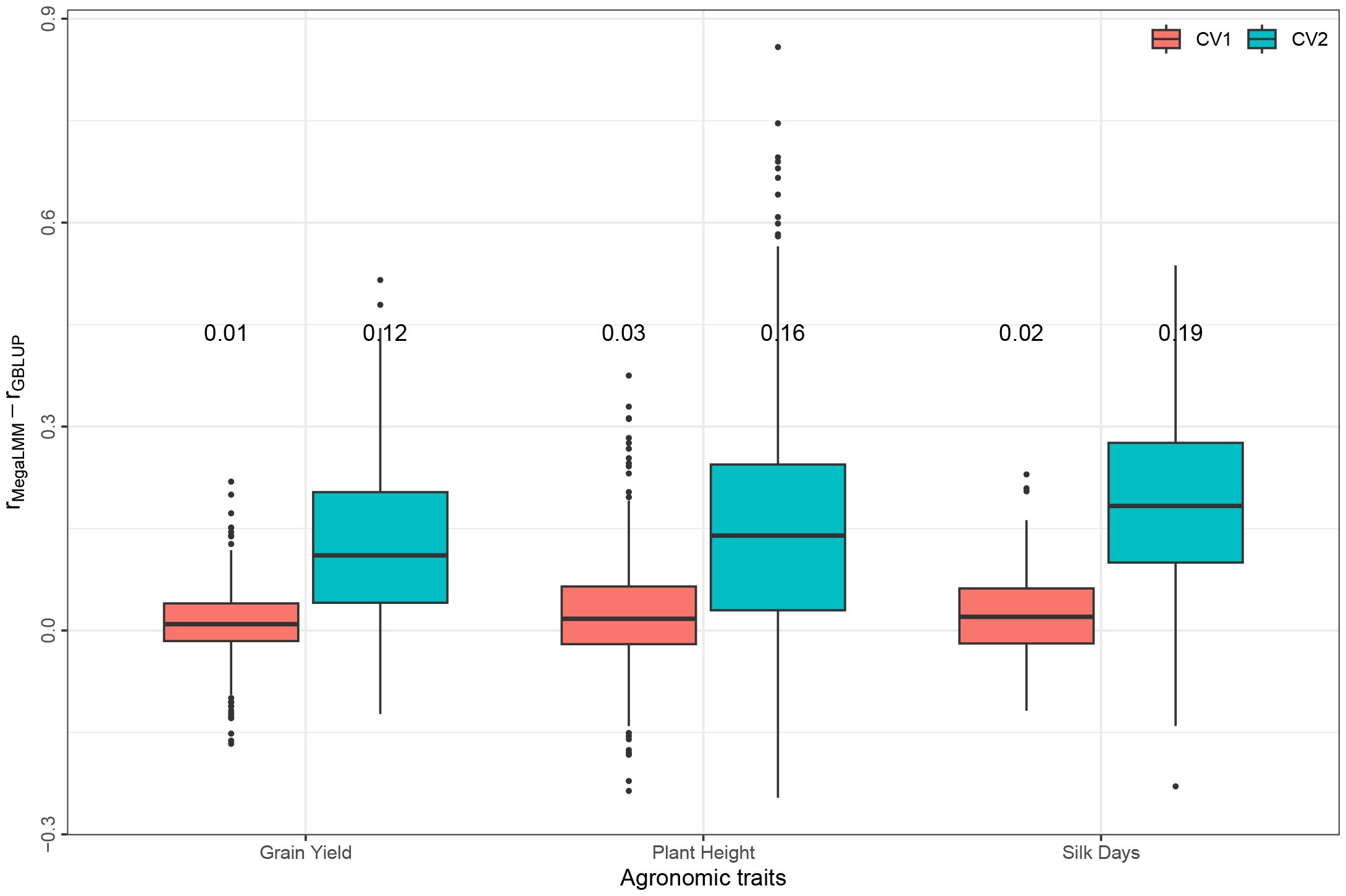
Predictive ability difference between MegaLMM and GBLUP for three agronomic traits (Silk Days, Plant Height, and Grain Yield). Each point within a boxplot represents the predictive ability difference between MegaLMM and GBLUP for a specific experiment. The mean predictive ability difference for each trait within each scenario is shown above the corresponding boxplot.

**Figure S2.**
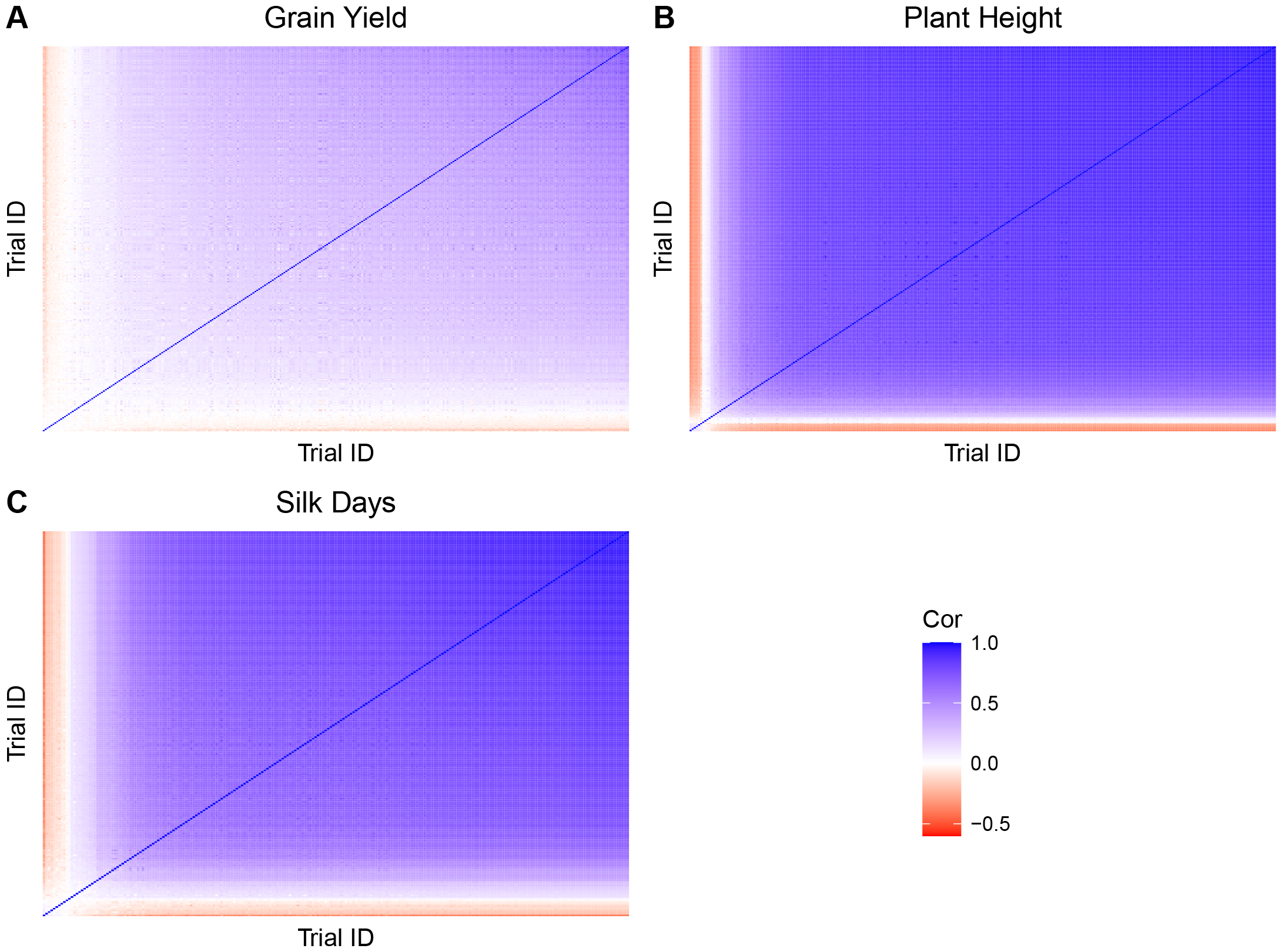
Pairwise phenotypic correlation between experiments estimated by MegaLMM for three agronomic traits (Silk Days, Plant Height, and Grain Yield)

**Table S1.** Description of the 11 weather environmental variables used in our study.

| Parameter | Units | Long Name/Description |
| --- | --- | --- |
| T2MWET | C | Wet Bulb Temperature at 2 Meters |
| QV2M | g/kg | Specific Humidity at 2 Meters |
| RH2M | % | Relative Humidity at 2 Meters |
| T2M_MAX | C | Temperature at 2 Meters Maximum |
| ALLSKY_SFC_SW_DWN | MJ/m<sup>2</sup>/day | All Sky Surface Shortwave Downward Irradiance |
| PS | kPa | Surface Pressure |
| T2MDEW | C | Dew /Frost Point at 2 Meters |
| WS2M | m/s | Wind Speed at 2 Meters |
| T2M_MIN | C | Temperature at 2 Meters Minimum |
| T2M | C | Temperature at 2 Meters |
| PRECTOTCORR | mm/day | Precipitation Corrected |

## Notes

### Competing Interest Statement

The authors have declared no competing interest.

